# Effect of Data Heterogeneity in Clinical MALDI-TOF Mass Spectra Profiles on Direct Antimicrobial Resistance Prediction through Machine Learning

**DOI:** 10.1101/2024.10.18.617592

**Authors:** Youngjun Park, Michael Weig, Christine Noll, Oliver Bader, Anne-Christin Hauschild

## Abstract

The matrix-assisted laser desorption-ionization time-of-flight mass spectrometry has become a powerful tool for accurate species identification in routine diagnostic microbiology. Recently, the application of machine learning models with MALDI-TOF mass spectra data indicated that rapid prediction of antimicrobial resistance patterns might facilitate even timelier and improved antimicrobial treatment. Although MALDI-TOF mass spectra data have proven valuable for clinical decision support, the issue of class imbalance in routine clinical data is often overlooked. This imbalance arises from factors such as local epidemiology, selective pressure from antibiotics, culture conditions, the methodology of phenotypic antimicrobial susceptibility testing, and sample preparation processes. Here, we provide a large mass spectra dataset, MS-UMG, for antimicrobial resistance prediction model training. With previously available public datasets, our dataset is evaluated and validated for usage in AMR prediction. We further explore the mass spectra data and identify informative regions on the spectra profile for AMR prediction. Moreover, we investigate the composition of this clinical dataset and present the implications of data heterogeneity on machine learning model performance. In conclusion, our findings highlight that accurate comprehension of clinical routine data and consideration of diverse hospital protocols are critical for effective clinical decision support systems with machine learning models.

**Key Points:**

- Introduced a large-scale clinical mass spectrometry dataset to the scientific community for research on antimicrobial resistance.
- Conducted a comparison and evaluation of this dataset with other existing large-scale MS datasets, highlighting its value for developing and validating predictive models in clinical settings.
- Demonstrated the robustness of machine learning models for antimicrobial resistance prediction using large-scale clinical mass spectra profiles.
- Analyzed the impact of data heterogeneity on the training and performance of machine learning models, emphasizing the need to account for variability in clinical routine data to enhance model reliability and generalizability.

## Introduction

Species identification of bacteria and fungi using traditional methods can be a time-consuming and complex task. Distinguishing species among phenotypically, biochemically, or genetically similar groups is particularly challenging [1]. In most clinical laboratories today, matrix-assisted laser desorption-ionization time-of-flight (MALDI-TOF) mass spectrometry (MS) has been established as the most reliable method for species identification over the past decade, virtually replacing all other phenotype-based tests [2]. Consequently, mass spectrometry (MS) data from these analyses is now being massively and continuously produced globally [3].

Previous studies indicate that MALDI-TOF MS data harbor numerous untapped biomarkers that can be utilized to further distinguish bacterial lineages below the species level [4, 5]. Additionally, these biomarkers may be correlated with specific phenotypic traits [6], including the presence of drug resistance mechanisms [7, 8, 9, 10, 11].

Clinical diagnostics also requires determining drug susceptibility profiles of clinical isolates to assess downstream therapeutic options and treatment or hygiene regimes [12, 13]. However, today, this is only possible to a certain degree by using the species information, e.g., guiding the choice for antibiotics active in Gramnegative or Gram-positive organisms [14] and using known intrinsic resistances of the identified species. Thus, subsequent phenotypic antimicrobial susceptibility testing (AST) is critical to provide valuable information about acquired resistance traits in clinical isolates. These AST data and established clinical breakpoints then enable therapeutic adjustments, such as high-dose therapy (in intermediate sensitive isolates) or the use of alternative anti-infective regimens (in resistant isolates). In addition, the early detection of multi-resistance in the inpatient setting is of immediate relevance for isolation measures in the context of hospital hygiene in order to prevent the spread of multi-resistant pathogens [15, 16]. However, these tests require additional time-consuming culture steps, where the bacteria are grown in the presence of different antimicrobial substances using reference broth microdilution, standardized disk diffusion, or automated AST [17, 18]. Depending on the laboratory setup, the applied AST method, and the growth characteristics of the isolate, these AST methods may delay targeted adjusted therapy by one or more days.

MALDI-TOF MS can effectively identify specific protein fingerprints that are associated with known antimicrobial-resistant clonal groups. An example of a successful application is the differentiation of *Bacteroides fragilis* strains carrying the *cfiA* metallo-betalactamase gene[19], the detection of biomarkers correlating with the presence of oxacillin resistance in *Staphylococcus aureus*, [20] or for plasmid-encoded resistances such as the KPC gene [9, 10, 11]. These findings suggest that MALDI-TOF MS data contains crucial information that can indicate the presence of antimicrobial resistance.

Building onto this foundation, a recent study by Weis *et al*. [13] demonstrated the suitability of machine learning (ML) for predicting antimicrobial drug resistance (AMR) with high precision and recall from MALDI-TOF MS data. Such approaches are highly desirable for implementation in clinical contexts, as resulting machine learning algorithms can be expected to profoundly impact time-to-first-result and thus improve early treatment decisions [21]. This usage of ML for predicting antimicrobial resistance based on clinical data shows considerable potential, yet significant challenges remain due to data imbalance and heterogeneity.

Data imbalance arises from clinical procedures such as sample preparation, culture techniques, and storage conditions, which can disproportionately favor certain microorganisms. Culture media plays a central role in microbiology laboratories for the isolation, cultivation, and identification of microorganisms. The components of these media can significantly influence which types of bacteria or other microbes will grow and thrive under specific conditions. This selective growth environment allows researchers to focus on particular organisms of interest while minimizing the presence of unwanted contaminants [22, 23]. Additionally, hospitals often focus on specific pathogens prevalent in their patient populations, resulting in an under-representation of rare pathogens or resistance patterns. Heterogeneity in clinical datasets is further exacerbated by the batch effects: different equipment, laboratory environment, and different levels of staff expertise across institutions [24]. Regional differences in pathogen prevalence and institutional priorities also contribute to the variability.

To develop robust and reliable machine learning models for antimicrobial resistance prediction, it is crucial to understand underlying constraints originating from clinical protocols [25]. Considering the representation of minority and majority classes is crucial when learning from imbalanced data. Models tend to be biased towards the majority class, often predicting it more frequently and ignoring the minority class. Additionally, high accuracy can be misleading in imbalanced datasets. Therefore, addressing class imbalance is essential to ensure fair and effective model performance across all classes [26].

In this study, we provide a new dataset of resistance information on antimicrobials and MALDI-TOF mass spectra of University Medical Center Göttingen (MS-UMG). To demonstrate the added value, we present a comparative analysis using various ML models for the prediction of antimicrobial resistance on the newly presented MS-UMG data and the DRIAMS-A-D datasets [13] as comparison sets. Most importantly, we investigated the impact of the previously described biases in training data from clinical routine processes on the training and evaluation of the ML models.

## Results

### MS-UMG dataset

We aggregated MALDI-TOF MS data of organisms isolated from clinical specimens from the University Medical Center Göttingen (UMG) in 2020 / 2021 during routine diagnostic procedures and integrated these with corresponding antimicrobial susceptibility profiles. In total, this amounted to 26,961 mass spectra and their corresponding metadata entries for the year 2020, and 50,381 mass spectra and their corresponding metadata entries for 2021, respectively. The dataset reflects 348 different species of bacterial and fungal organisms and 72 different antimicrobial susceptibility testing (AST) results. To allow for compatibility and straightforward integration with the DRIAMS A-D datasets data was processed and organized into the same structure as previously described [13]. Detailed information about data pre-processing is outlined in the Methods section. The MS-UMG dataset is publicly available at https://zenodo.org/records/13693800.

### Validation of the MS-UMG dataset in a machine learning model to predict antimicrobial resistance

Three ML models were used to assess antimicrobial resistance prediction using the MS-UMG dataset. We demonstrated the clinical utility of routine MS data for predicting antimicrobial resistance with Logistic Regression (LR), a tree-based model (LightGBM), and multilayer perceptron (MLP) on three important pathogens with well-understood resistance mechanisms, namely (*Escherichia coli, Klebsiella pneumoniae*, and *Staphylococcus aureus*). As the main AMR prediction scenarios, we tested *E. coli* with LightGBM, *K. pneu-moniae* with MLP, and *S. aureus* with LightGBM (Figure 1). These scenarios were chosen for comparison with previous work by Weis *et al*. [13].

**Figure 1.**
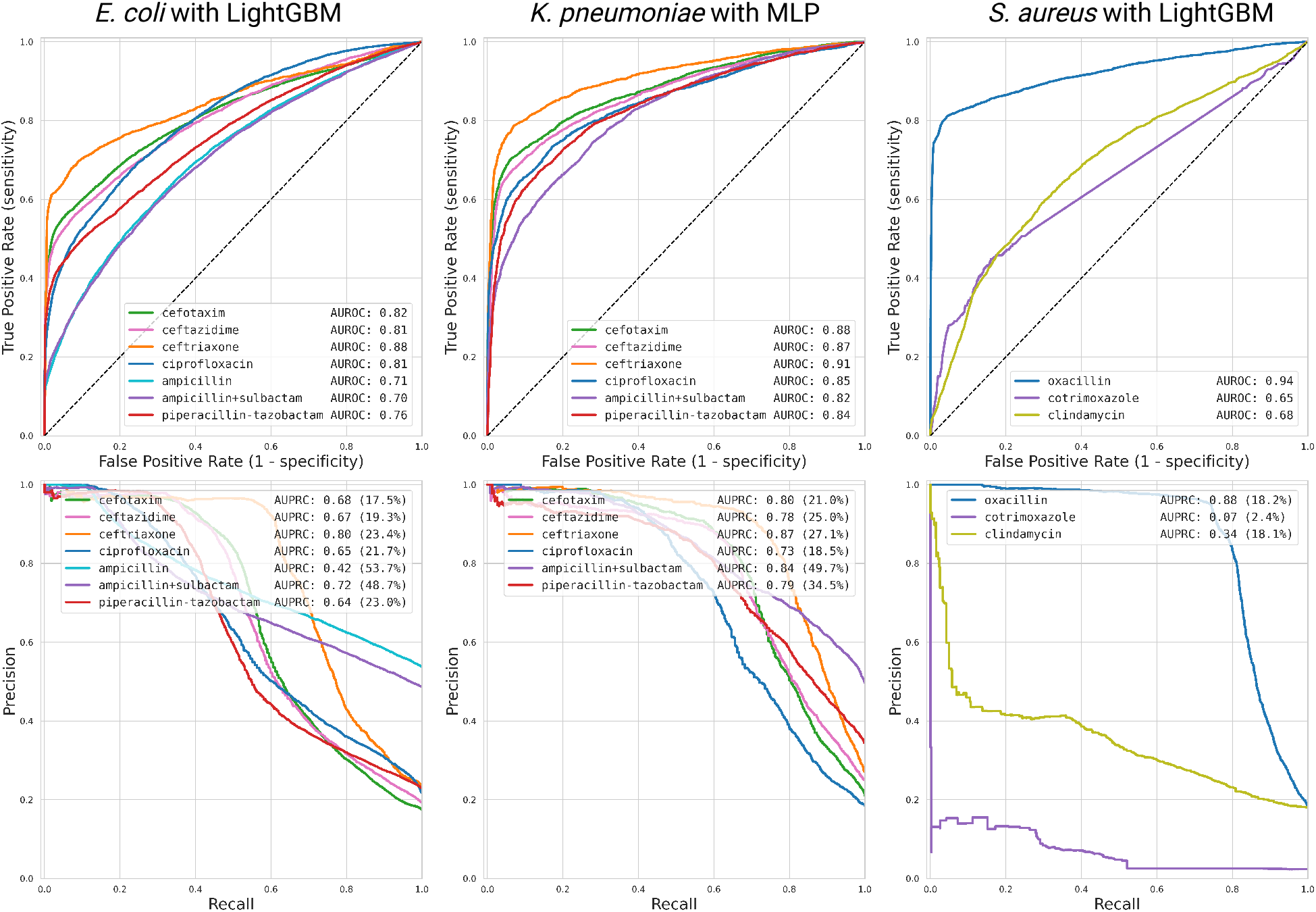
Receiver operating characteristic and precision-recall curves of AMR prediction models on MS-UMG. Three machine-learning model results are presented. Left column: *E. coli* (LightGBM), middle column *K. pneumoniae* (MLP), right column *S. aureus* (LightGBM).

The models exhibited high overall performance in both AUROC and AUPRC metrics for both *E. coli* and *K. pneumoniae*, indicating that the classifiers can provide accurate antimicrobial resistance predictions. Using oxacillin resistance as a measure in *S. aureus*, MLP similarly predicted MRSA phenotypes at high AUROC and AUPRC. However, when used on Cotrimoxazole or Clindamycin measures, both AUROC and AUPRC remained low (AUROC=0.65 / AUPRC=0.07, and AUROC=0.68 / AUPRC=0.34, respectively). This may likely be due to the fact that the training data does not discriminate underlying resistance mechanisms, which are more heterogenic in these cases: Resistance to Cotrimoxazole can be based on the acquisition of genes from the auxiliary genome [27], mediating changes of the permeability barrier, the transcription of efflux pumps or target enzymes, or mutations of the target enzymes [28]. Similarly, in the case of Clindamycin (Lincosamides), resistance can be due to a variety of target-site modifications, increased gene expression of efflux pumps, or enzymatic drug inactivation [29].

### Mass range analysis for information content

The original study that shared the first large-scale dataset utilized the full spectrum profile range of 2,000–20,000 Da for AMR prediction [13]. They reported that informative feature bins related to resistance were characterized by having a mass-to-charge ratio (m/z) value of less than 10,000 Da. However, no further analysis was conducted to assess the information density of the spectral profiles. Therefore, we examined the ideal bins range for predicting antimicrobial resistance from mass spectra peaks: First, we divided the spectra into 1,500 Da ranges and trained ML models on them. We analyzed three microbe-antibiotic pairs, as presented in the previous results, using the DRIAMS-A and MS-UMG datasets. Figure 2 shows the trend in AUROCs of the machine learning models trained on different bin ranges. The ML models trained on spectra from the lower mass ranges demonstrated superior predictive performance as compared to those trained on spectra from the higher mass ranges, suggesting that the lower mass ranges contain more relevant or informative features for accurately predicting AMR. Conversely, the higher range spectra might include more noise or less distinctive markers, leading to a decrease in model accuracy. This trend was clearly observable in both datasets (Figure 2).

**Figure 2.**
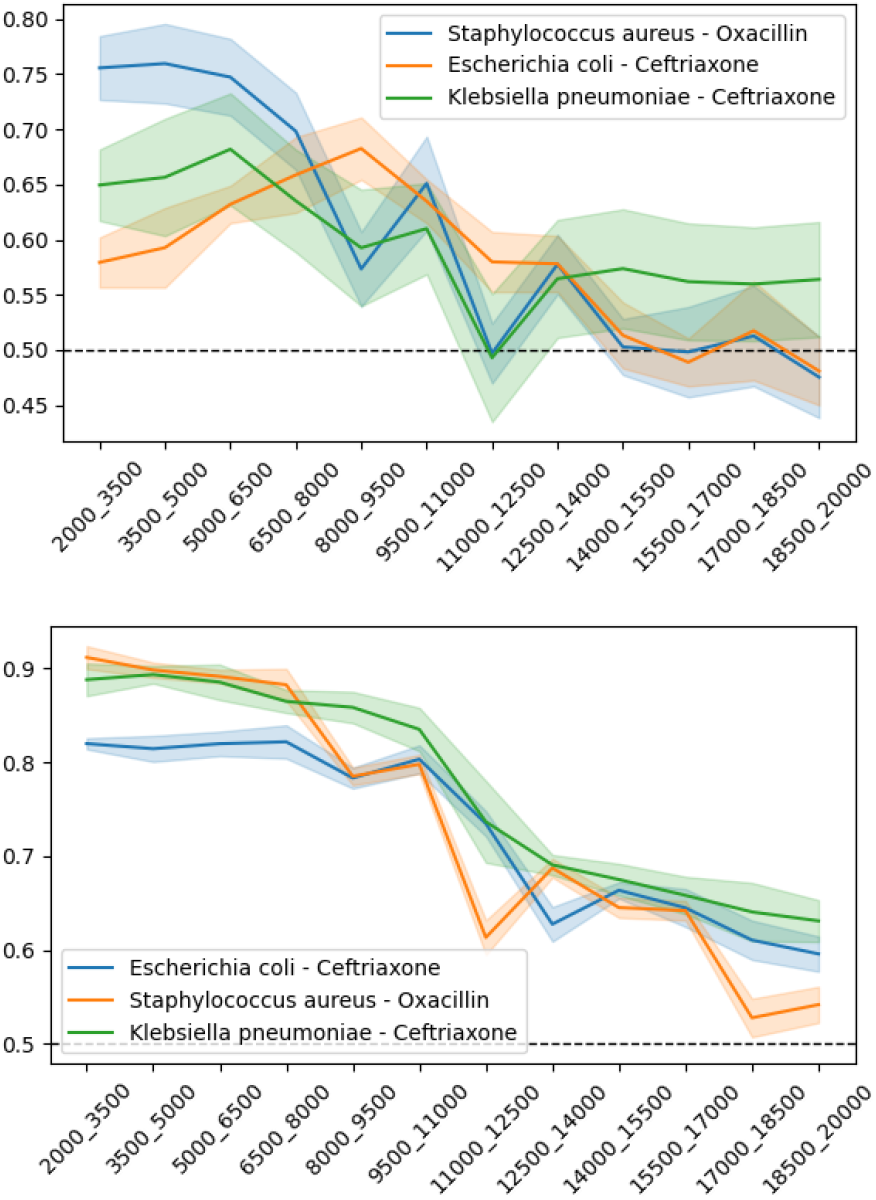
AUROC of AMR prediction with range of 1500Da spectra profile. Three microbes and antibiotic pairs are investigated for informative spectra ranges. The machine learning models, LightGBM for *E. Coli* and *S. aureus*, MLP for *K. pneumonia*, are trained with a subset of spectra bins. Both DRIAMS-A (above) and MS-UMG (below) datasets were analyzed, and the plots indicate that the lower Da range plays a more significant role compared to the higher Da range.

Next, we, therefore, trained models using spectra accumulated from the lower bound, starting at 2,000 Da (Figure 3a), and compared their performance with a respective model trained on spectra accumulated from the upper bound, starting at 20,000 Da (Figure 3b). Progressively incorporating bins containing spectral peak information from the lower and upper bounds, assessed how the inclusion of different spectral ranges affected the respective model’s predictive abilities. Here, we observed that the lower range of mass spectra contained more concentrated information relevant to AMR prediction. Notably, models trained using only the 2,000–5,000 Da range achieved predictive performance comparable to those utilizing the entire mass range. This finding suggests that critical biomarkers or features associated with AMR are primarily localized within this lower mass range, making it a more efficient choice for predictive modeling. This aligns with observations that more intensive signals are observed in this range than in the 10-20 kDa range, which is intrinsic to the limitation of the measurements [30, 31].

**Figure 3.**
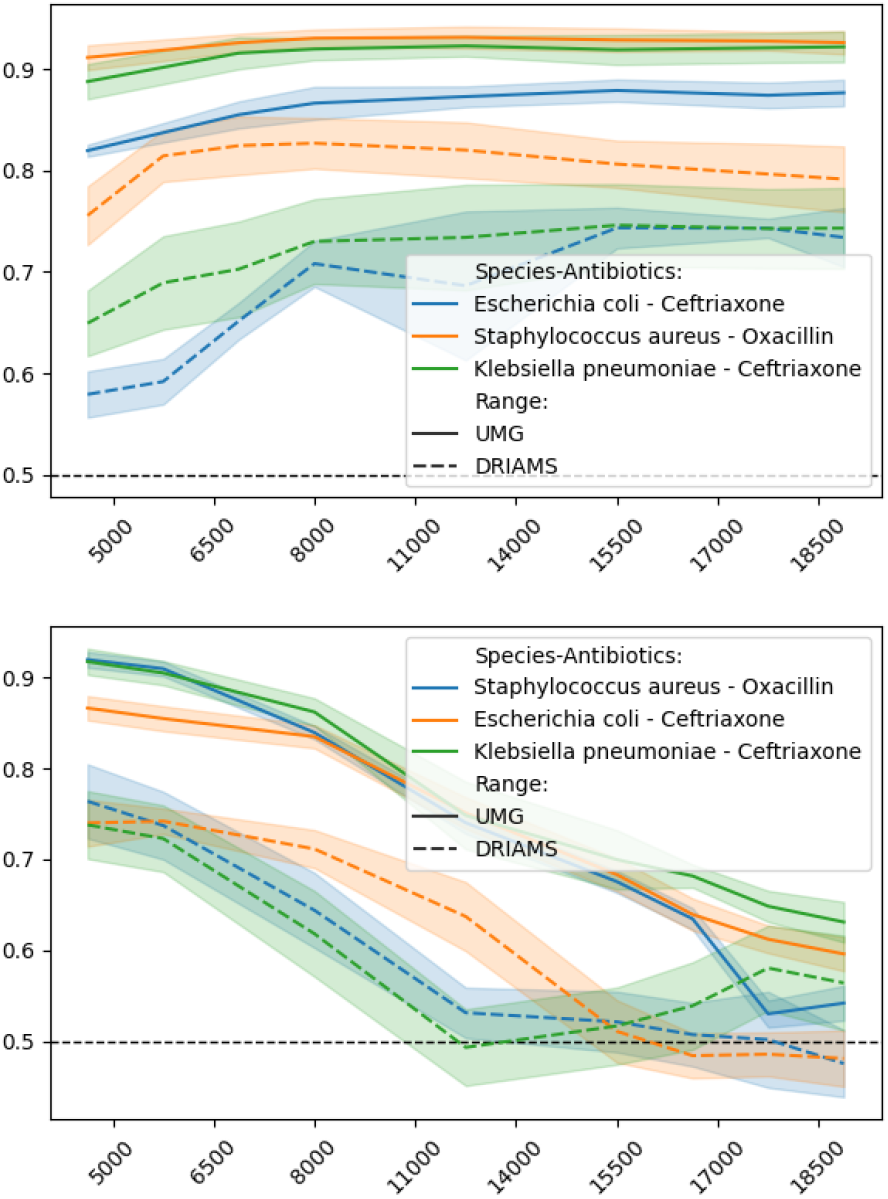
Examining the AUROC values obtained when different spectra range from the lower bound and the higher bound used for training. Three microbes and antibiotic pairs are investigated for informative spectra ranges. The machine learning models, LightGBM for *E. Coli* and *S. aureus*, MLP for *K. pneumonia*, are trained with a subset of spectra ranges. Both DRIAMS-A (above) and MS-UMG (below) datasets were analyzed. Plots show that the AUROC scores from both the lower and higher mass-to-charge ranges crossed within the 5000-6500 Da region. The higher Da range spectra show a declining AUROC, implying that these regions contribute less to the AMR prediction.

### Data bias originated in clinical routine process deteriorate machine learning model

The ML models demonstrated significant potential in AMR prediction as a clinical decision support system. The predictive performance of the described ML models on the MS-UMG dataset shows higher AUROC and AUPRC values compared to those from the DRIAMS-A dataset. These results might indicate higher homogeneity of MS-UMG data compared to the DRIAMS dataset (Figure A3). Next, we aimed to identify the factors contributing to this enhanced predictive performance. Clinical routine MS data for the UMG-MS data set data was obtained from cultures using two major workflows: regular agar (“regular”) and a hygiene screening pipeline using chromogenic media (“screening”). The latter employs antibiotic-containing culture media, and thus specifically enriches for community-derived AMR strains.

To identify the impact of data from screening agar on the ML model, we further divided the MS-UMG dataset along these constraints. We then compared the performance of AMR prediction using the same ML models on these distinct subsets. This comparison allowed us to evaluate how the different culture methods influenced the model’s ability to predict antimicrobial resistance. Indeed, the model trained exclusively with screening agar data achieved (4) higher AUROC and AUPRC as compared to both other models trained solely with regular agar data or the combined datasets. This suggests that the biased nature of the screening agar samples towards resistant phenotypes enhances the model’s predictive performance for AMR. The screening agar dataset exhibited a highly imbalanced class ratio. For instance, in the *E. coli*-Ceftriaxone and *S. aureus*-Oxacillin scenarios, the screening agar data contained 99% resistant phenotypes and only 1% susceptible phenotypes. Retrospective inspection of the 1% of cases in the “S” screening category showed that they likely represented false negative data from AST, e.g. a case of inducible expression without the proper tests performed as evidence for this phenotype was below thresholds. Interestingly, this case was predicted as “R” by the ML model trained on regular agar data. This case further indicates the potential of the ML model as a complementary method to AST.

### Comparison of MS-UMG to DRIAMS dataset

In all three main scenarios analyzed, removing the screening agar data from the dataset led to a decrease in the performance of AMR prediction (Figure. The results obtained from the model trained exclusively on the regular agar dataset became comparable to the results by Weis *et al*. [13]. This result shows the significant impact of screening agar data on the ML model training by inflating the model’s performance.

For a more detailed comparison of our MS-UMG data with the DRIAMS datasets, we performed cross-site validation of machine learning models. We evaluated whether the predictive performance achieved with MS-UMG data was transferable to other DRIAMS sites, and vice versa. MS-UMG and DRIAMS-A–D datasets were divided into training and test datasets. We then trained models lightGBM models on *E. coli* and *S. aureus* and MLP on *K. pneumoniae* on each site, and tested across all sites. The ML model trained on the MS-UMG dataset demonstrated comparable predictive perfor-mance when applied to other datasets within the DRIAMS collection. Particularly, the LightGBM model for *E. coli* on Ceftriaxone had an AUROC of 0.764 on DRIAMS-B. The MLP model for *K. pneumoniae* on Ceftriaxone displayed an AUROC of 0.704 on DRIAMS-C, and LightGBM model for *S. aureus* on Oxacillin an AUROC of 0.671 on DRIAMS-B (Figure A3).

After excluding the screening sample data, the overall performance of the model on the MS-UMG dataset declined. However, the cross-dataset performance between the DRIAMS dataset and the MS-UMG dataset improved (Figure 5). This suggests that while the model’s ability to perform well on the MS-UMG dataset alone was reduced, its capacity to generalize across different datasets was enhanced. Notably, the model trained exclusively on regular agar displayed better generalization, indicating that it captured more fundamental or generalizable features.

**Figure 4.**
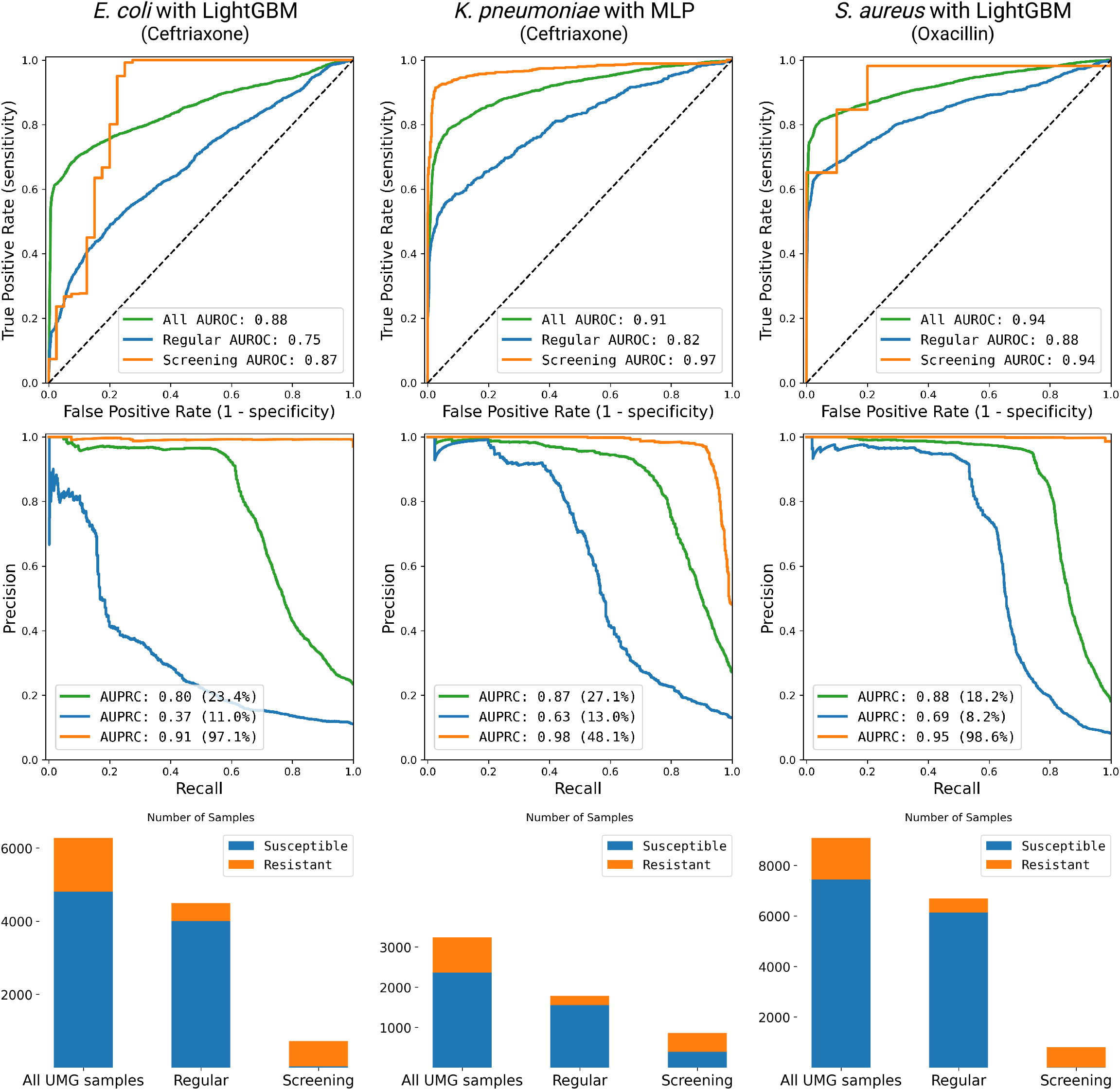
Receiver operating characteristics of accuracy and precision and class distribution in samples from regular and screening agar. The three main scenarios, E-CEF, K-CEF, and S-OXA, are presented. The differences in antimicrobial resistance prediction are shown by varying training dataset compositions. Bar plots present class distribution in each dataset of regular and screening agar.

**Figure 5.**
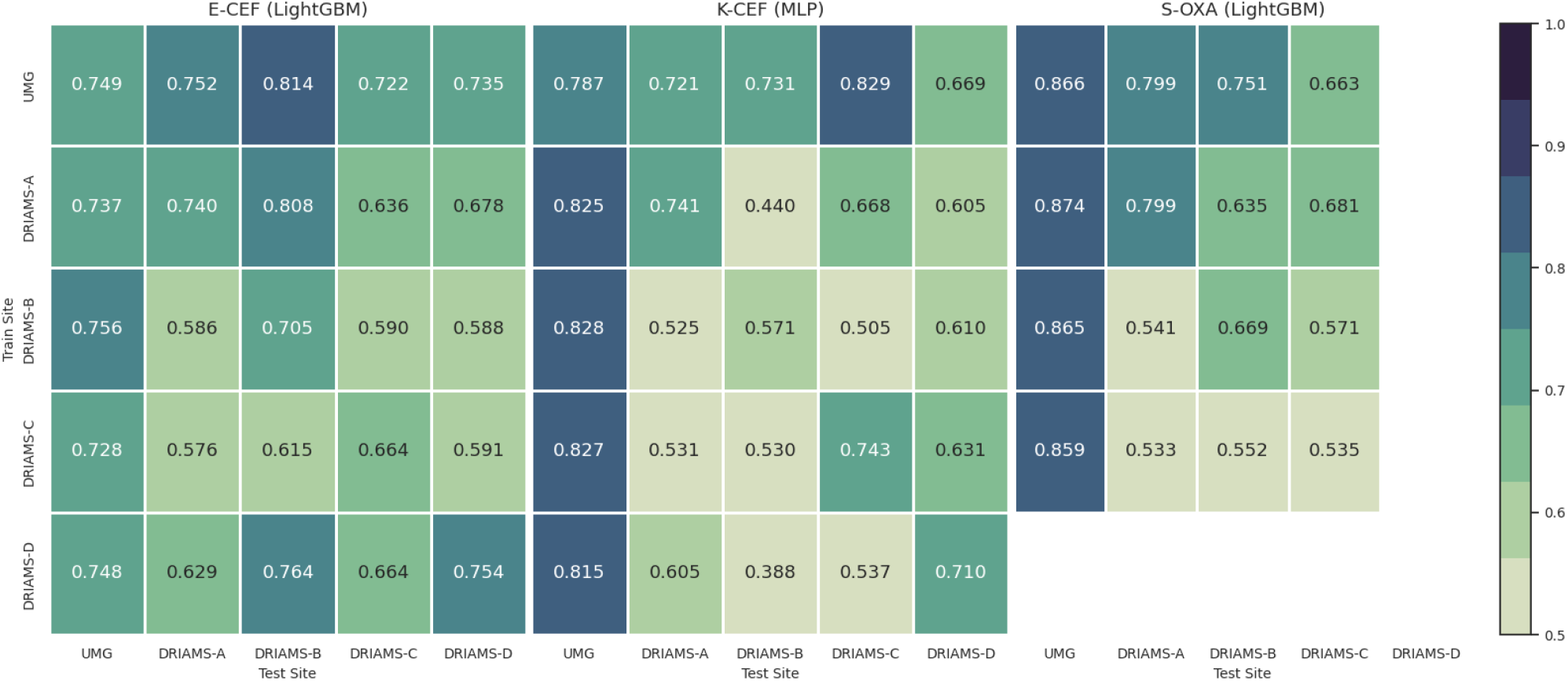
Validation of predictive performance on different sites using regular agar data. Heatmap shows the mean AUROC performance of different pairs of train-test datasets in three main scenarios of antimicrobial resistance prediction. E-CEF is *E. coli* with Ceftriaxone, K-CEF is *K. pneumoniae* with Ceftriaxone, and S-OXA is *S. aureus* with Oxacillin. Each analysis is 5-fold cross-validated with fixed random seed and the AUROC number is averaged of 10 repeats. The data of DRIAMS pairs are derived from the original work.

### Feature importance analysis reveals impact of data bias

To investigate further the impact of biased samples from the screening system on the prediction of antimicrobial resistance using a machine learning model, we analyzed the importance of features with Shapley value analysis.

As data from screening media inflated the performance of AMR prediction in ML models, the feature importance within these models must be also influenced by this data heterogeneity. For comparison, the Shapley values were calculated for the ML models trained on three subsets of the MS-UMG dataset: regular, screening, and combined data. This analysis is able to show that the predictor appears to base its two positive class predictions (resistant/intermediate) on two key factors: either the presence of high-intensity values, indicated by red, or the absence of any measured intensity, shown as blue. We focused on the analysis of *S. aureus* with Oxacillin and the 20 important bins from three differentially trained models are presented (Figure 6).

**Figure 6.**
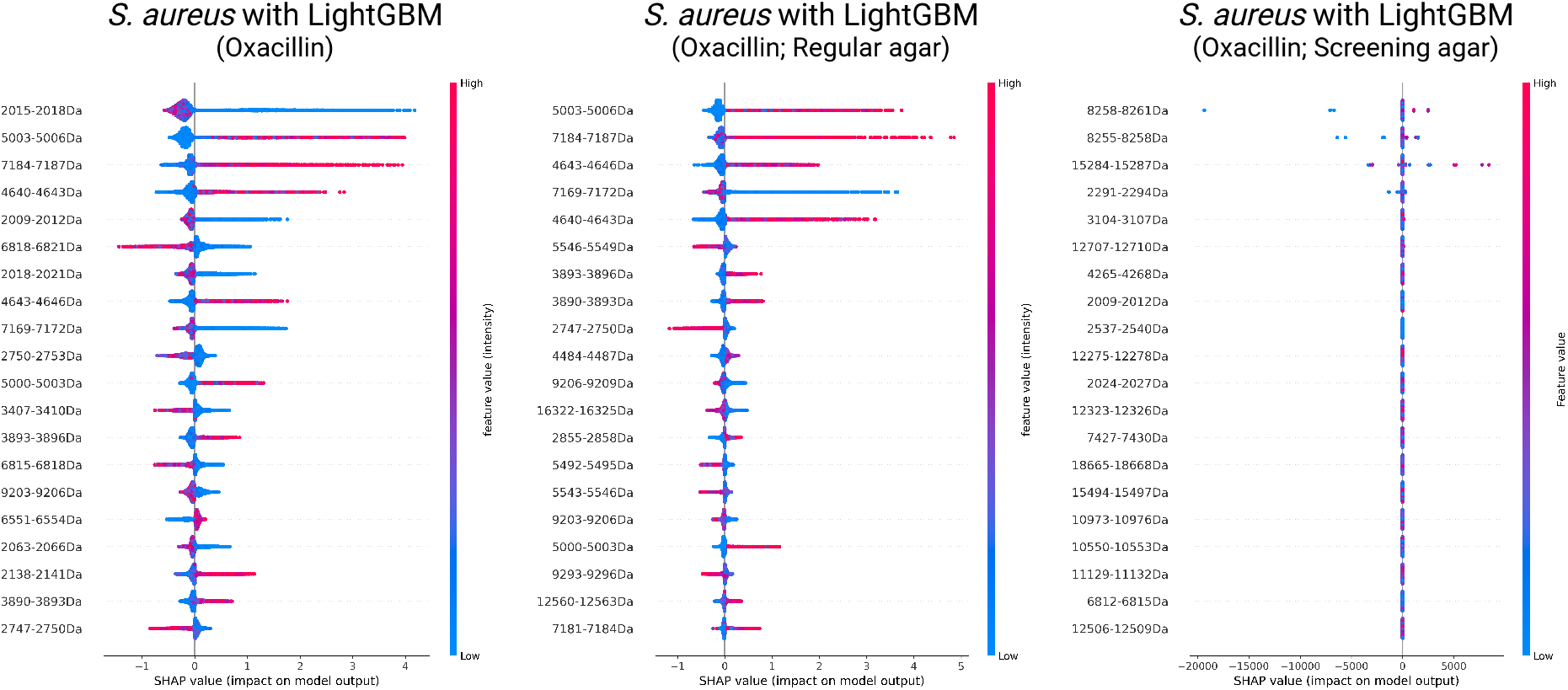
Shapley analysis on the S-OXA scenario with a LightGBM model. The left plot presents the Shapley values when all samples are used for machine learning analysis. The center plot shows the Shapley values when only regular agar samples are used for the machine learning analysis. The right plot shows the Shapley values when only screening agar samples are used for the machine learning analysis.

In the top 20 important bins, the bin ranges 3890–3893, 4640–4643, and 5003–5006 are identified, which are consistent with the findings from the DRIAMS study [13]. Particularly, the feature importance ranking of the bin range 3890–3893 is higher in the model trained without the inclusion of screening agar. When screening agar is included in the training data, the model tends to focus on different bins, potentially overlooking those related to proteins associated with oxacillin resistance. The analysis suggests that the bin range 3890–3893 may contain information relevant to a protein with a molecular mass of 3,891 Da, a mass attributed to the uncharacterized *S. aureus* protein SA2420.1. This interesting finding from our comparisons of differentially trained models is related to already reported protein related to differentiating between methicillin-susceptible *S. aureus* (MSSA) and methicillin-resistant *S. aureus* (MRSA), as well as distinguishing between various sublineages within MRSA itself.

Similarly, the Shapley analyses of *E. coli* and *K. pneumoniae* with Ceftriaxone also showed a change in rank among important bins (Appendix Figure A1, A2). Notably, the Shapley analysis of *E. coli* with screening agar identified 9737-9740 as important bins. This is consistent with previous findings reporting a 9,738 Da peak in extended-spectrum beta-lactamases (ESBL) negative and a 9,736 Da peak in ESBL-positive *E. coli* [32]. ESBL production is the most common mechanism of ceftriaxone resistance in *E. coli*[33].

## Discussion

In this work, we introduce a new large-scale MALDI-TOF MS dataset to the research community. We envision that this dataset could facilitate innovative research, ultimately leading to more effective strategies for machine learning-based infection diagnosis and antibiotic optimization, which are crucial in the global fight against antibiotic resistance. Furthermore, to the best of our knowledge, this is the first report on the effect of data heterogeneity in antimicrobial resistance prediction using MALDI-TOF mass spectrometry data via machine learning models.

The application of machine learning models to predict antimicrobial resistance using the MS-UMG dataset demonstrates the clinical utility of routine mass spectrometry data in combination with machine learning for AMR prediction. This approach aligns with the broader trend of leveraging machine learning and deep learning models for the detection and prediction of AMR, as highlighted in recent works [13]. MALDI-TOF coupled with ML techniques has been used successfully to identify genetically similar species, to detect antibiotic resistance or virulence factors, and to detect epidemic clones [34, 35, 13]. This new machine learning-based approach has great potential in clinical decision support systems to speed up and optimize antimicrobial therapy.

Data imbalance can affect machine learning models in several ways. Firstly, models trained on imbalanced data may become biased towards the majority class, leading to poor performance in minority classes. This bias can result in inaccurate predictions that fail to identify rare but clinically significant cases. Secondly, the model may overfit the majority class, failing to generalize well to new, unseen data, undermining its effectiveness in real-world applications. Finally, the model may have reduced sensitivity in detecting rare but potentially critical resistance patterns, which is a significant concern in clinical diagnostics where detecting such patterns is crucial for patient care [26].

While MALDI-TOF mass spectrometry combined with machine learning offers great potential for antimicrobial resistance prediction, the challenge of data imbalance must be carefully addressed. By implementing appropriate strategies to handle imbalanced data and leveraging large, well-curated datasets, we can develop more accurate and reliable models for clinical decision support. This approach will ultimately lead to improved patient care and more effective antimicrobial stewardship in healthcare settings.

## Materials and methods

### MS-UMG data acquisition

The MS-UMG dataset connects raw mass spectrometry data with susceptibility testing data, obtained from May 2020 to June 2021 under routine diagnostic conditions. The MS data was generated on two individual routine diagnostic machines, a Microflex LT-SH and a SMRT System (both Bruker Daltonics, Bremen, Germany), using the MALDI Biotyper Software suite (V4.1, database version 16). Susceptibility data was obtained in parallel to mass spectrometry using the Vitek2 system (Biomerieux, Marcy l’Etoile, France), and resulted in either ‘S’ (susceptible), ‘I’ (susceptible, increased exposure), or ‘R’ (resistant) values for each substance measured according to the EUCAST interpretative breakpoint guidelines. MS profiles and susceptibility data were mapped using the internal numbering of the laboratory information system, and subsequently fully anonymized with UUID.

### Mass spectrometry data processing

The spectra are extracted from the BioTyper systems in the Bruker Flex data format and processed with the MALDIquant Package v1.19 in R. Mass spectra profiles are converted to 6,000 lengths vectors with bin size of 3 Da across all analyses. This bin size was chosen for direct comparability of the MS-UMG to the DRIAMS datasets [13]. The vector’s intensities of each spectrum were transformed using the square root method and are smoothed with the SavitzkyGolay method with a half window size of 10. The baseline is removed in 100 iterations with the SNIP algorithm. The intensity is then calibrated using the total ion current (TIC), and lastly, the spectra are trimmed to the range of 2,000 20,000 Da.

MS-UMG data is further dissected into two different subgroups, regular and screening agar, and analyzed. The regular agar dataset contains 24,833 samples for the year 2020 and 33,216 samples for the year 2021. The screening agar dataset contains 2128 samples for the year 2020 and 3595 samples for the year 2021.

### Public dataset

These contain mass spectra and antimicrobial susceptibility profiles collected as part of the clinical routine from a set of Swiss hospitals, with different time frames: DRIAMS-A, November 2015–August 2018; DRIAMS-B, January 2018–June 2018; DRIAMS-C, January 2018–August 2018; and DRIAMS-D, January 2018–June 2018 [13]. The publicly available real-world clinical microbiology datasets are integrated and analyzed together with our MS-UMG dataset.

### Machine learning analysis

We first compared ML algorithms for the prediction of antimicrobial resistance on the MS-UMG dataset in a similar scheme to the work from Weis et al. [13]. These were logistic regression (LR), LightGBM [36], and multi-layer perceptron (MLP). Parameters are optimized through grid search with 5-fold cross-validation. For reproducibility, a random seed number is given as an argument, and it controls the train/test data split.

For the ML analysis, the intensity measurements were binned using a window size of 3 Da to establish a fixed dimensionality for the input features. The m/z axis having the range from 2,000 to 20,000 Da, is dissected into equal-sized, 3 Da, bins and sums the intensity of all peaks in the same bin. All the patient samples’ MS data were preprocessed with the same procedure as previous work [13]. The preprocessed 6,000 binned vectors were used for the machine learning training and testing.

Our primary performance metric for the evaluation is AUROC. Given the substantial class imbalance in the datasets for most antibiotics (both DRIAMS and MS-UMG datasets), AUROC proves advantageous as it remains unaffected by the class ratio, allowing for comparability across antibiotics with varying class ratios. However, relying solely on AUROC may be misleading as it does not reflect precision. AUPRC was also provided as a supplement evaluation metric to address this limitation. The majority of the performance metrics presented in figures and tables represent the mean performance derived from a five-fold cross-validation. In instances of significantly limited training data during five-fold cross-validation, the approach is adjusted to three-fold cross-validation. Shaded areas refer to the standard deviation across the respective evaluation metric.

Three ML models are employed to predict AMR using large-scale clinical routine MS data. In this work, we evaluated the performance of LR and LightGBM. MLP to predict antibiotic resistance from mass spectra features of patient samples. The baseline pipeline used in this work is maldi-learn library developed by Weis *et al*. (available in https://github.com/BorgwardtLab/maldi-learn) [13]. It has a set of three ML models for classification for AMR prediction: LR, LightGBM, and MLP. For the implementation of LR and MLP, the scikit-learn package in Python is used. For the LightGBM, the official implementation of the LightGBM package by Microsoft Corporation is used [36]. The entire pipeline is wrapped using pipeline of scikit-learn package. Analysis code and results used in this work are available in https://gitlab.gwdg.de/cdss/maldi-tof_ms_amr_umg

The analysis follows the following order: The default setup for cross-validation was 5-fold cross-validation. Thus, samples were randomly split into a training and test dataset with an 80% to 20% ratio, while stratifying for the AMR phenotype and the species. The hyperparameters were optimized through grid search by optimizing AUROC with the training data. This grid search included different penalties (L1, L2, no penalty), scaling methods (standardization or none), and regularization parameters (1e-3, 1e-2, 1e-1, 1e0, 1e1, 1e2, 1e3). With the best hyperparameters, the model was retrained with the full training dataset, predicted the test dataset, and evaluated.

## Acknowledgement

Y.P., O.B., and AC.H. conceived this project. M.W., C.N., and O.B. collected and aggregated clinical routine MALDI-TOF mass spectra and antimicrobial resistance screening profiles. Y.P. performed data processing and machine learning analysis. All authors interpreted the results and prepared the manuscript.

## Appendix

**Figure A1.**
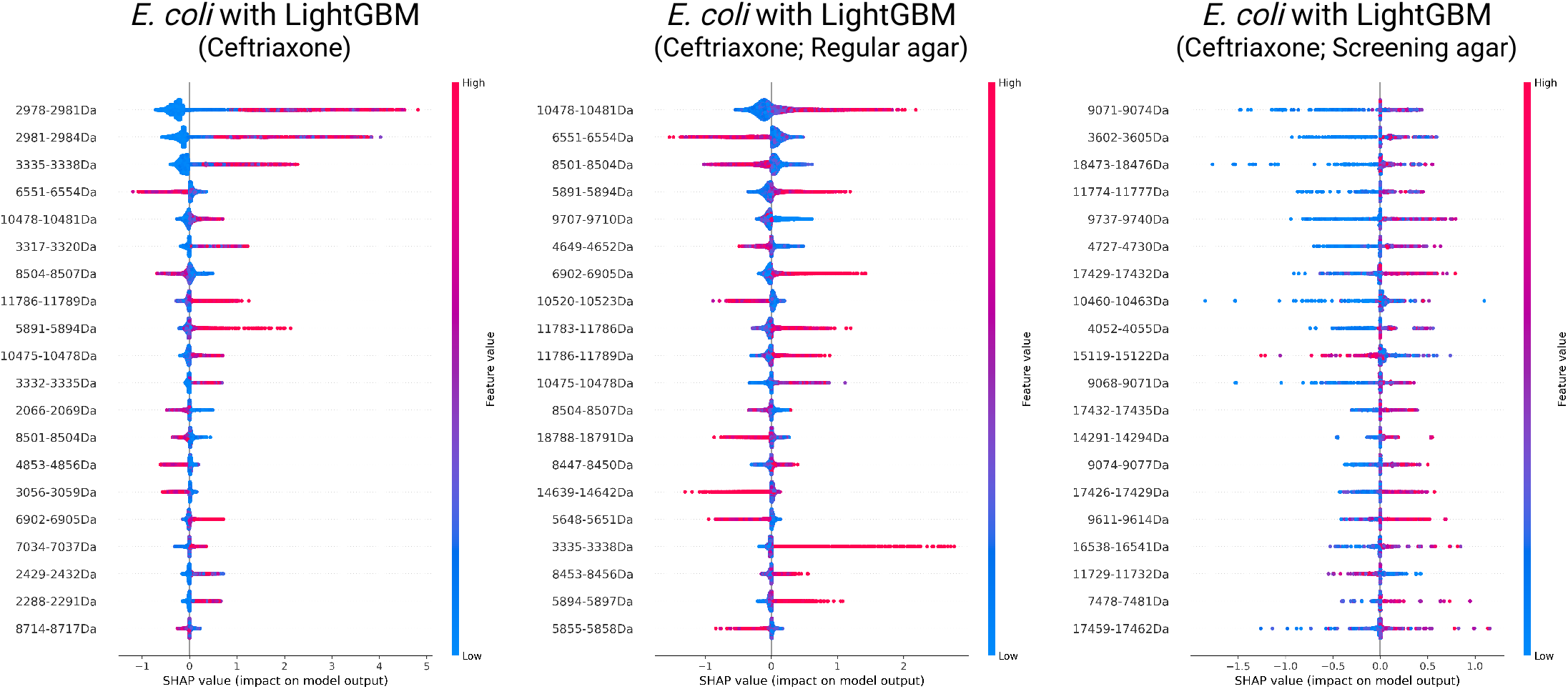
Shapley analysis on the E-CEF scenario with a LightGBM model. The left plot presents the Shapley values when all samples are used for machine learning analysis. The center plot shows the Shapley values when only regular agar samples are used for the machine learning analysis. The right plot shows the Shapley values when only screening agar samples are used for the machine learning analysis.

**Figure A2.**
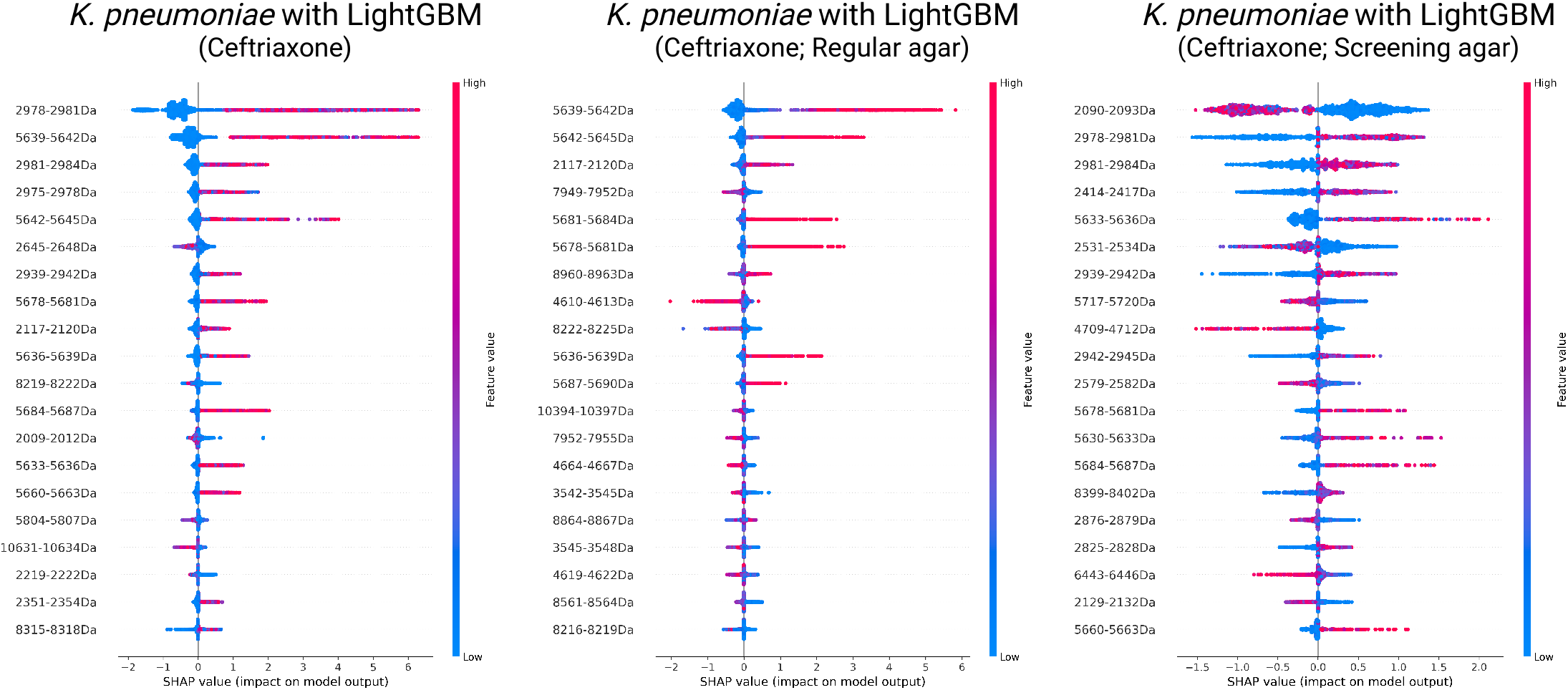
Shapley analysis on the K-CEF scenario with a LightGBM model. The left plot presents the Shapley values when all samples are used for machine learning analysis. The center plot shows the Shapley values when only regular agar samples are used for the machine learning analysis. The right plot shows the Shapley values when only screening agar samples are used for the machine learning analysis.

**Figure A3.**
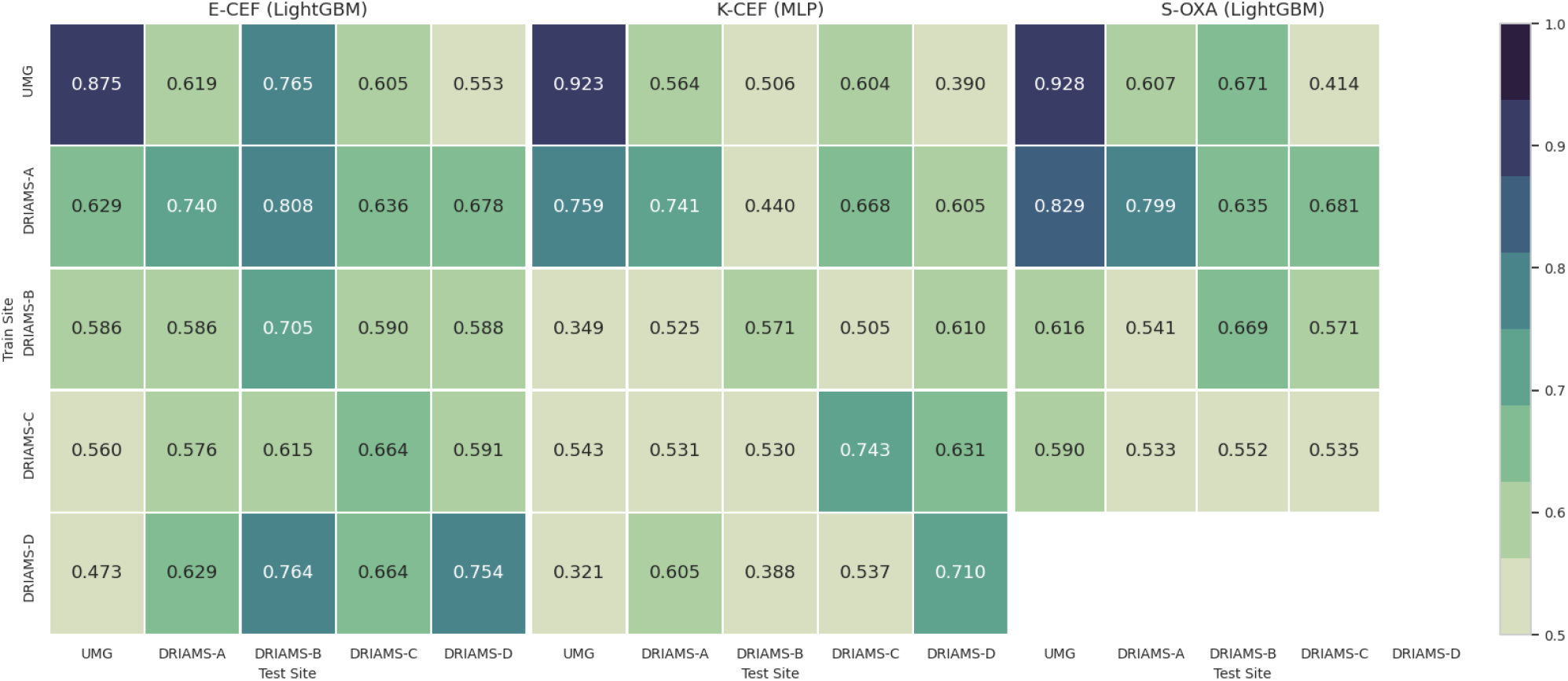
Validation of predictive performance on different sites using entire MS-UMG data. Heatmap shows the mean AUROC performance of different pairs of train-test datasets in three main scenarios of antimicrobial resistance prediction. E-CEF is *E. coli* with Ceftriaxone, K-CEF is *K. pneumoniae* with Ceftriaxone, and S-OXA is *S. aureus* with Oxacillin. Each analysis is 5-fold cross-validated with fixed random seed and the AUROC number is averaged of 10 repeats. Thus, the training dataset and test dataset is same set in all the analyses. The data of DRIAMS pairs are derived from the original work.

## Notes

https://zenodo.org/records/13911744

### Competing Interest Statement

The authors have declared no competing interest.

